# Upstream trophic structure modulates downstream community dynamics via resource subsidies

**DOI:** 10.1101/088096

**Authors:** Eric Harvey, Isabelle Gounand, Chelsea Little, Emanuel A. Fronhofer, Florian Altermatt

**Affiliations:** Department of Evolutionary Biology and Environmental Studies, University of Zurich, Winterthurerstrasse 190, CH-8057 Zürich, Switzerland; Eawag, Swiss Federal Institute of Aquatic Science and Technology, Department of Aquatic Ecology, Überlandstrasse 133, CH-8600 Dübendorf, Switzerland

**Keywords:** meta-ecosystems, directional flows, cross-ecosystem subsidies, river ecosystems

## Abstract

In many natural systems, the physical structure of the landscape dictates the flow of resources. Despite mounting evidence that communities’ dynamics can be indirectly coupled by reciprocal among-ecosystem resource flows, our understanding of how directional resource flows might indirectly link biological communities is limited. We here propose that differences in community structure upstream should lead to different downstream dynamics, even in the absence of dispersal. We report an experimental test of the effect of upstream community structure on downstream community dynamics in a simplified but highly controlled setting, using protist microcosms. We implemented directional flows of resources, without dispersal, from a standard resource pool into upstream communities of contrasting interaction structure and then to further downstream communities of either one or two trophic levels. Our results demonstrate that different types of species interactions in upstream habitats may lead to different population sizes and levels of biomass in these upstream habitats. This, in turn, leads to varying levels of detritus transfer (dead biomass) to the downstream communities, thus influencing their population densities and trophic interactions in predictable ways. Our results suggest that the structure of species interactions in directionally structured ecosystems can be a key mediator of alterations to downstream habitats. Alterations to upstream habitats can thus cascade down to downstream communities, even without dispersal.

## Introduction

In many natural systems, the physical structure of the landscape dictates the flow of organisms and resources. Previous work has shown that directionally biased movement of organisms can have significant effects on species co-existence (Levine 2003, Lutscher et al. 2005, 2007, Salomon et al. 2010), meta-population dynamics (Fronhofer and Altermatt 2015) and stability (Elkin et al. 2008), and meta-community structure (Altermatt et al. 2010, Dong et al. 2016, Bourgeois et al. 2016). Mounting evidence now suggests that communities’ dynamics can be indirectly coupled by the reciprocal spatial exchange of resources, even in the absence of dispersal (Loreau et al. 2003, Gravel et al. 2010b, Harvey et al. 2016). Yet, attempts to look at effects of directionally biased movement of resources on community dynamics are scarce (but see Polis and Hurd 1995 on detrital inputs from sea to islands), and contrary to research on reciprocal exchanges (Leibold et al. 2004, Gravel et al. 2010a, Gounand et al. 2014), there is no general understanding of how directional resource flows might indirectly link biological communities.

The understanding of directional resource flows and how they might link biological communities in space is especially relevant for ecosystems or communities in which resource flows are dictated by gravity or dominant wind patterns, such as river ecosystems, mountain slope habitats or vertically structured plant communities. For example, the “river continuum concept” (Vannote et al. 1980) suggests that shifts in local community structure along river branches are the sole result of linearly changing physical conditions, and that downstream communities profit from upstream energy processing inefficiencies. A direct but yet unexplored implication of such a linear transfer of energy is that differences in community structure upstream should lead to different downstream dynamics, even in the absence of dispersal: because biotic interactions modify the way energy is distributed among the different species, the interaction structure of an upstream community should determine the quality and quantity of resources (e.g., dead cells from various species with contrasting stoichiometry and inorganic resources from metabolic waste) flowing through to downstream communities. Therefore, all else being equal, the same amount of resources assimilated by different upstream communities may lead to the production of qualitatively very different subsidies (Gounand et al. in review). In a system with reciprocal subsidy exchanges this could alter source-sink dynamics (Gravel et al. 2010a) or nutrient co-limitation where communities exchange different limiting resources (Marleau et al. 2015). However, in ecosystems with strong directionality, upstream communities are likely to act as mediator of the effects of resource flow on downstream communities (Fig. 1a).

**Figure 1.**
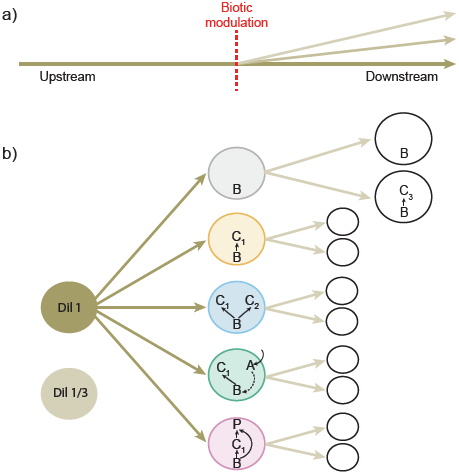
a) In many ecosystems, resource flow is directionally biased; as they move downstream, these resources will be integrated, processed and modified by biotic communities (biotic modulation) meet along the way with potentially important implications for downstream community dynamics. In our experiment (b), starting from an initial resource pool (brown circle: standard protist medium, either none-diluted or one-third diluted), we test the effect of contrasting upstream community structures (descending order from trop: *bacteria alone, monoculture, competition, facilitation*, and *predation*) on bacteria populations in two downstream communities with different trophic structures (one vs. two trophic levels). The two first downstream communities are enlarged to exemplify composition and internal dynamics; analogue settings were present for all downstream systems. B: mixture of three bacteria species (*Serratia fonticola, Bacillus subtilis* and *Brevibacillus brevis*), C_1_: *Colpidium sp.*, C_2_: *Paramecium aurelia*, A: *Euglena gracilis*, P: *Daphnia pulicaria*, C_3_: *Tetrahymena pyriformis*.

As a first demonstration, we here report an experimental test of the effect of upstream community structure on downstream community dynamics in a simplified but highly controlled setting. Using protist microcosms, we implemented directional flows of nutrients moving from a standard resource pool into upstream communities of contrasting interaction structure (“*Monoculture*”, “*Competition*”, “*Predation*”, “*Facilitation*”, “*Bacteria alone*”, see Fig. 1b), and then to further downstream communities of either one (bacteria) or two trophic levels (bacteria and a consumer, Fig. 1b). We tracked population densities of bacteria and protists in the downstream communities and linked them to the respective upstream community structure.

## Methods

We studied the effects of directional spatial flows mediated by biotic modulation in sequentially linked communities (called either “upstream” or “downstream”, corresponding to the flow direction, Fig. 1). We manipulated the structural composition of the upstream community and monitored subsequent effects on the downstream community in the absence of dispersal (i.e., only spatial flows of resources).

To test the effect of upstream community structure on downstream community dynamics we built a factorial protist microcosm experiment composed of 10 types of two-patch meta-ecosystems linked by directional spatial flows. Each two-patch meta-ecosystem was composed of an upstream community, which was either bacteria alone (a mixture of *Serratia fonticola, Bacillus subtilis* and *Brevibacillus brevis*, referred to as the “*Bacteria alone*” treatment), the same bacteria mixture and the bacteriophagous ciliate *Colpidium* sp. (“*Monoculture*”), the same bacteria mixture and *Colpidium* sp. with the bacteriophagous *Paramecium aurelia* (“*Competition*”), or with the autotroph *Euglena gracilis* (“*Facilitation*”, see Fig.1b), or with the generalist predator *Daphnia pulicaria* (“*Predation*”, see Fig.1b). As our focus is on the effect of different upstream community structures on downstream community dynamics, we use only the treatment rather than species names in the text for the sake of clarity and consistency. The choice of each species combination is based on prior knowledge from previous experiments in similar settings (Gounand et al. in review, Carrara et al. 2015, Harvey et al. 2016). These five upstream communities were either connected to a downstream community composed of bacteria alone (one-trophic-level community) or bacteria with the bacteriophagous *Tetrahymena pyriformis* as consumer (two-trophic-level community). To test the sensitivity of our results to initial resource concentration and thus the generality of our findings on the effects of upstream community structure on downstream dynamics, we also replicated our experiment with two different initial inflowing resource levels (Fig.1b). To do this we either did or did not dilute by one third the standard protist medium (Carolina Biological Supply, Burlington NC, USA, 0.46 g protist pellets 1 L^−1^ tap water) that was added to the upstream community twice a week (see “Diffusion” section below, and Fig.1b). Each of the ten two-patch meta-ecosystems was replicated four times for a total of 160 microcosms.

Each microcosm consisted of a 250 mL Schott bottle that was filled to 100 mL. Microcosms were assembled by first adding 75 mL of pre-autoclaved and filtered (Whatman filters) standard protist medium, and 5 mL of bacteria inoculum. After 24 hours, to allow time for bacteria growth, we added 20 mL of protist culture with each protist species at carrying capacity (10 mL per species for mixed communities, 20 mL of *Colpidium* sp. for *Monoculture* communities, and 20 mL of *Tetrahymena pyriformis* for the two-trophic-level downstream communities). Thus, protist communities were added at 20% of their carrying capacity and were allowed to grow 24 hours before the first resource flow event, henceforth referred to as diffusion and described below. In upstream communities with predation, we added 5 individuals of *Daphnia pulicaria* in each microcosm. For further details on general methods used in our protist microcosm experiments, see Altermatt et al. (2015).

### Diffusion

The directional flow of resources from upstream to downstream communities was carried out in three distinct steps to ensure the maintenance of a constant volume in each microcosm. First, 30 mL was removed from each downstream community. Second, 30 mL from each upstream community was sampled and microwaved to turn all living cells into detritus (Harvey et al. 2016). After a 3 hours cooling period at ambient temperature (20 °C), the microwaved samples had reached 20 °C and were poured into the respective downstream recipient ecosystems. Third, 30 mL of autoclaved standard protist medium (none diluted or 1/3 diluted according to treatment) was added to each upstream microcosm from the same homogenized medium pool to ensure that effects to downstream communities were not caused by differences in intake resource quality. This manipulation resulted in a directed resource flow from the common resource to the upstream community, and from the upstream community to the downstream community (Fig. 1b).

Because our main focus was on the mediating effect of upstream community structure on downstream communities via resource flows only, we chose microwaving until boiling as a method to kill living cells, ensuring that no dispersal could occur between our microcosms. While small molecules are likely lysed during boiling, we cannot exclude that other substance than nutrients, potentially acting as kairomones are diffused.

Previous work showed that chemical cues from live or dead con- and heterospecifics can be used to inform movement and dispersal decisions (Hauzy et al. 2007, Fronhofer et al. 2015b, 2015a) with important consequences for population growth and large-scale spatial dynamics (Fronhofer et al. in press). However, our main conclusions on the effects of upstream community structure on downstream ecosystem dynamics are consistent with expectations from previous work on nutrient flow effects in similar settings (Harvey et al. 2016). Therefore we are confident that a majority of the effects we find are due to flows of nutrients. Based on previous work in similar experimental settings, we also know that 30% diffusion represents the best trade-off to maximize effects of spatial flows while minimizing the mortality effect associated with the procedure (Gounand et al. in review, Harvey et al. 2016).

### Measurements

Measurements were synchronized with diffusion events. The measurements occurred every Monday and Thursday (experimental days 0, 3, 7, 10, and 14 respectively), and diffusion occurred every Tuesday and Friday. At each measurement day, two 0.5 mL aliquots were sampled for each microcosm: one for protist and one for bacteria density analysis. Protist density was measured by using a standardized video recording and analysis procedure (Pennekamp and Schtickzelle 2013, Pennekamp et al. 2015). In short, a constant volume (17.6 μL) of each 0.5 mL aliquot was measured under a dissecting microscope connected to a camera and a computer for the recording of videos (5 sec./video, see Appendix S1 in Online Supporting Information for further details on this method). Then, using the R-package bemovi (Pennekamp et al. 2015), we used an image processing software (ImageJ, National Institute of Health, USA) to extract the number of moving organisms per video frame along with a suite of different traits for each occurrence (e.g., speed, shape, size) that could then be used to filter out background movement noise (e.g., particles from the medium) and to identify species in a mixture (see Appendix S1). Finally, for bacteria we measured densities using standard flow cytometry on fresh SYBR green fixated cells using a BD Accuri™ C6 cell counter (1/1000 dilution, following protocols in Altermatt et al. 2015).

### Statistical analyses

We analyzed effects of directed resource diffusion on downstream population dynamics of bacteria and *Tetrayhmena* separately. To test for the effect of upstream community on downstream community dynamics we used a three-way Linear Mixed Effect model (LME) testing the interactive influence of upstream community structure, presence of a second trophic level in downstream community, and continuous time on log-transformed bacteria density in the downstream communities. In parallel, for *Tetrahymena* (in the two-trophic-level downstream communities) we performed a two-way LME testing for the interactive effects of upstream community structure and continuous time on log transformed densities. In both models, to control for temporal pseudo-replication issues we added replicates and time as nested random factors.

Because we were also interested in linking changes in downstream communities to changes in upstream communities, as a complementary analysis, we also tested differences in bacteria and protist densities among the different upstream community structure treatments. To this end, we used a two-way LME testing for the interactive effects of upstream community structure and continuous time on log-transformed densities. We also added replicates and time as nested random factors.

For each LME model, we used an AIC-based simplification procedure, removing terms sequentially, starting with the highest level of interactions. While we fitted models during model selection using maximum likelihood (“ML”) and the “BFGS” optimization method (Nash 1990), the final models were refitted by maximizing the restricted log-likelihood (“REML”, see Pinheiro et al. 2016). We used standardized residuals versus fitted-value plots, residual distribution, variance over-dispersion, and log-likelihood information to select the most appropriate transformation for each model. Finally, even if there was not always significant variations over time (i.e., Fig. 2b), for the sake of clarity and consistency, we extracted predictions for each LME over time along with 95% confidence intervals, which we report here as our main results (Fig.2). We interpreted treatments with none-overlapping confidence intervals as significantly different. As complementary information, between treatment differences are reported as mean ± standard deviation in text. The statistical model tables can be readily reproduced by using the provided R-script (Dryad – DOI: dox.doi.org/XXXXXX).

**Figure 2.**
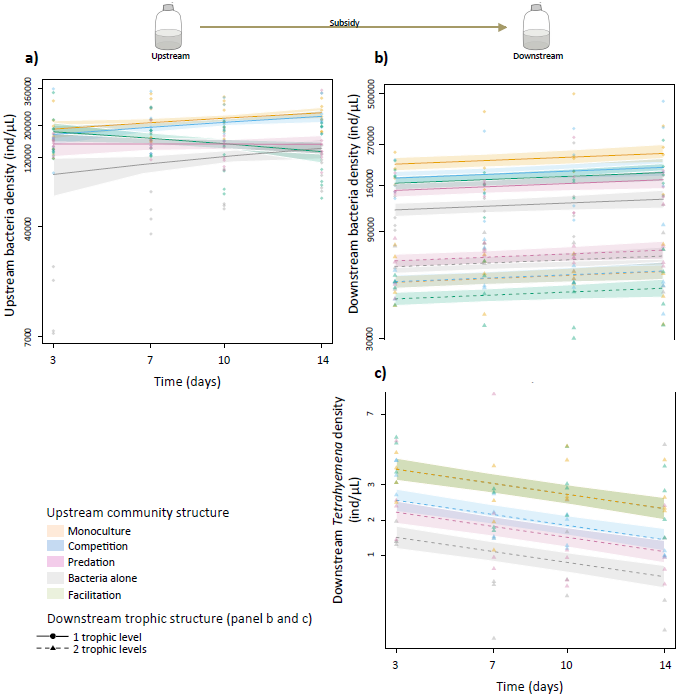
Effect of upstream community structure on upstream bacteria density (panel A), and on downstream bacteria (panel B) and *Tetrahymena* (panel C) densities in the one-trophic-level (full lines – *Tetrahymena* absent) and in the two-trophic-level (dashed lines – *Tetrahymena* present) communities. Points (*Tetrahymena* asbent) and triangles (*Tetrahymena* present) represent raw data. Full and dashed lines represent model predictions with 95% confidence intervals as shadings. Y-axes on all panels are on log-scale, but for clarity tick numbers represent raw densities. On panel C, model predictions for *Facilitation* (2.81 ± 1.25 ind/μL) and *Monoculture* (2.76 ± 1.05 ind/μL) are completely overlapped, and on panel B (dashed lines) *Monoculture* (55370 ± 15486 ind/μL) is visible just under *Competition* (55670 ± 17169 ind/μL).

All analyses were conducted with R 3.1.2 (R Core Team 2016), using the ‘bemovi’ package (Pennekamp et al. 2015) for video analyses, the “nlme” package for statistical modelling (Pinheiro et al. 2016), and the “car” (Fox and Weisberg 2011) and “MASS” (Venables and Ripley 2002) packages to identify proper variable transformations for statistical analyses. All data and the main r-script to reproduce the results can be downloaded from Dryad (DOI: dox.doi.org/XXXXXX).

## Results

We examined how downstream communities of varying trophic structure (one or two trophic levels) are influenced by the structure of interactions in upstream communities (see Fig.1b). Changing the dilution factor of the resources flowing through upstream communities led to no qualitative differences in community dynamics, however in the diluted treatments population densities of bacteria and *Tetrahymena* were overall two to five times lower and the range in densities were greatly dampened (for bacteria; between 1400 and 240000 ind./μL in the diluted treatment and between 7400 and 500000 ind./μL in the none-diluted treatment, for *Tetrahymena*; between 0.003 to 1.10 ind./μL in the diluted treatment and between 0.3 and 10 ind./μL in the none-diluted treatment, see Fig.S1 for paralleled low-dilution treatment results). Because the results were qualitatively the same, we will further report results only for the none-diluted treatment (but see Fig.S1). As expected, we found that the main driving factor of bacteria populations in the downstream communities was the local presence of a second trophic level (here *Tetrahymena*), which greatly reduced bacteria densities and dampened variation in population densities across treatments regardless of the upstream community structure (from 186000 ± 76000 to 58000 ± 18000 ind./μL, see Fig.2b). However, when looking at each downstream community separately we found significant influences of the upstream communities on bacteria dynamics coupled through resource flows only.

More specifically, in the one trophic level downstream communities, bacteria density was highest when the upstream community is a *Monoculture* (240000 ± 96000 ind./μL) or a *Competition* (208000 ± 99000 ind./μL, Fig.2b) community, and was lowest when it contained only bacteria (*Bacteria alone*; 135000 ± 46000 ind./μL, Fig. 2b). These patterns matched with bacteria densities in the upstream communities where densities were consistently higher in *Monoculture* (216000 ± 60000 ind./μL, Fig.2a) and *Competitive* (202000 ± 66000 ind./μL, Fig.2a) communities and consistently lower in *Bacteria alone* communities (115000 ± 77000 ind./μL, Fig. 2a). In the upstream *Facilitation* communities, bacteria densities declined over time (Fig.2a) following the expected increase of consumer densities in this treatment (see Fig. S2 for *Colpidium* densities in upstream communities). The average bacteria density of 181000 ± 37000 ind./μL placed this treatment between the highest (*Monoculture* and *Competition*), and the lowest (*Predation* and *Bacteria alone*) treatments, in terms of density, which matched with the pattern observed for bacteria downstream where *Facilitation* also represented the average median density (Fig.2b). In summary, different community structures in upstream habitats supported different levels of bacteria densities, which in turn led to varying levels of detritus transfer (dead bacteria biomass) to the downstream communities, thus influencing their bacteria densities (Fig. 2ab).

In the presence of bacterivorous *Tetrahymena* (see Fig.2b), however, bacteria densities did not follow this consistent pattern: highest densities were instead found when the upstream community contained bacteria only (*Bacteria alone*; 66000 ± 16000 ind./μL, Fig.2b) or a *Predator* (71000 ± 19000 ind./μL, Fig.2b) and lowest when there was *Facilitation* (45000 ± 14000 ind./μL, Fig. 2b). Bacteria density patterns, in these two-trophic-level downstream communities, seem to match local *Tetrahymena* densities with highest bacteria densities found at lowest *Tetrahymena* densities (*Bacteria alone* and *Predation*, Fig. 2c) and lowest bacteria densities found at highest *Tetrahymena* densities (*Facilitation*, Fig. 2c). Instead of bacteria density (as observed in the one-trophic-level treatment), it is downstream *Tetrahymena* densities that match with bacteria densities in upstream communities with highest densities found for *Monoculture* (*Tetrahymena*: 2.76 ± 1.05 ind./μL, Fig.2c) and lowest densities found for *Bacteria alone* (*Tetrahymena*: 1.25 ± 1.07 ind./μL, Fig.2c). These results suggest varying levels of top-down pressure from *Tetrahymena* on bacteria in downstream communities as a function of varying upstream community structures.

## Discussion

We experimentally showed that upstream community structure affects downstream community dynamics through resource flows only. Our results demonstrate that different community structures support different bacteria densities in upstream habitats, which in turn lead to varying levels of detritus transfer (dead biomass) to the downstream communities, thus influencing their population densities and trophic interactions in predictable ways. In natural communities with many more species interacting, it is likely that different upstream community structures will also lead to qualitative changes in subsidy depending on the biomass distribution and the respective stoichiometric ratio of each trophic level (Gounand et al. in review, Sitters et al. 2015, Marleau et al. 2015). Overall, our work highlights that upstream communities can mediate the effect of resources flow on downstream communities, even in the absence of dispersal.

The presence of a consumer (here *Tetrahymena*) in the downstream communities greatly reduced prey (bacteria) density (see Figure 2). This top-down pressure varied as a function of upstream community structure: when bacteria density was higher upstream, there were more consumers downstream and less prey, suggesting a spatial cascade through subsidy. Therefore, it seems that the highest trophic level is the most sensitive to changes in resource flow. This result suggests that top-predators might be key to the response of local communities to variations in subsidy.

In upstream communities, monoculture and competition treatments had highest levels of bacteria compared to predation and facilitation. Low bacteria density in the predation treatment can be explained by our use of a large generalist predator (*Daphnia pulicaria*) that feeds both on bacteria and protists. In the facilitation treatment, the presence of the autotroph *Euglena gracilis* brings-in new resource through photosynthesis that benefits bacteria and likely increases grazing top-down pressure via a bottom-up trophic cascade – a pattern that we indeed observed in our results: decreasing bacteria density through time (Fig.2a), paralleled by an increase in the bacteriophagous *Colpidium* sp. (Fig. S2). These changes to resource quality and quantity may have cascaded to the downstream ecosystems. For instance, we know from previous work with *Daphnia* in similar settings that biomass tends to accumulate at the predator level and thus lead to a decline in detritus quality (increased in recalcitrant chitin content, see Gounand et al. in review), with negative consequences for connected ecosystems (Gounand et al. in review).Also, the presence of *Euglena* (facilitation treatment) generates a local enrichment effect, albeit limited by this species slow growth rate (Harvey et al. 2016). Overall, based on our results, downstream community dynamics seem to be mainly driven by variations in upstream bacteria densities (changes to subsidy quantity due to local upstream species dynamics), which likely acted in parallel to changes in resource quality and quantity from other internal dynamics in upstream ecosystems linked to our various community structure treatments (i.e., decreased quality in the *Predation* treatment and increased quantity in the *Facilitation* treatment). Our results thus clarify how upstream communities might affect outflowing subsidy quality/quantity and then cascade spatially to downstream communities. This study also emphasized that measuring detrital content should be a particular concern of future study to further elucidating specific mechanisms.

Interestingly in our upstream ecosystems we observed that bacteria densities were highest when growing with a consumer (*Colpidium sp.*), and lowest when growing alone. Despite that this result does not affect our main conclusion that pertains to the matching patterns between upstream and downstream communities as a function of upstream community structure, it is nonetheless a puzzling observation. Although we can only speculate on this, few hypotheses can however explain this counter-intuitive result (e.g., selective feeding). Because our bacteria community was composed of three species with wide interspecific size variations, selective feeding by the consumer, releasing one bacteria species from competition, thus leading to higher cell density (but not total biomass), appear to be most likely explanation.

Many natural ecosystems are characterized by directionally biased spatial flows of organisms and resources, such as alpine slopes, seashore habitats, vertical structure of tree or plant habitats, and river ecosystems, with the latter likely being the most studied. As opposed to many terrestrial systems, strong directional movements along dendritic-shaped networks dominate spatial processes in rivers (Altermatt 2013). These two fundamental attributes of river landscapes (directionality and dendritic-shaped network) have profound implications for the spatial distribution of diversity and local population dynamics (Carrara et al. 2012, Kuglerová et al. 2014, Seymour et al. 2015, Vitorino Júnior et al. 2016, Fronhofer and Altermatt in review). For instance, the river continuum concept (Vannote et al. 1980) suggests that specific communities form in rivers as a function of stream order (i.e., distance to the upstream source). These communities could not be maintained elsewhere because they require the specific physical conditions provided by their location in the river network (Vannote 1980) and recruitment from the directional movement of different upstream organisms from converging paths along the dendritic network (Muneepeerakul et al. 2008, Carrara et al. 2012). Despite the need for more empirical studies in different ecosystems to identify potential contingencies, our experimental results demonstrate that upstream community structure can act as a biotic modulator of resources thus indirectly affecting downstream community dynamics (Fig.1a), with important implications for landscape management and the mitigation of eutrophication issues in downstream habitats.

Our results suggest that upstream species interaction networks might be a key mediator of alterations to downstream habitats in directionally structured ecosystems. For instance, the impact of large nutrient loads from agricultural source upstream on downstream lakes could potentially be mitigated or amplified depending on the interaction structure of upstream communities. In our study we showed that a specific upstream community structure has the same qualitative effect on downstream dynamics regardless of initial resource concentration (none-diluted resource; Fig. 2 vs. diluted resource; Fig. S1). Our experiment thus suggests that biotic interactions *per se* might be a key mediator of spatial changes in community dynamics by indirectly linking communities via directional nutrient flows. This has significant, but yet untested implications for landscape management and the restoration of ecosystem services in ecosystems with directionally biased resource flows.

## Acknowledgements

We thank S. Gut and E. Keller for help during the laboratory work. Funding is from the Swiss National Science Foundation Grant PP00P3_150698.

## Supplementary Material

## Appendix S1 protist density counts by video analysis

The 5-second videos were recorded with a Canon camera, adjusted on a Nikon microscope with a 20-fold magnification, on 17.6μL-volume samples.

To process the videos, we first detected the moving particles with the functions locate_and_measure_particles(), link_particles() and filter_data() of the R-package bemovi (Pennekamp et al. 2015), coupled to the image analysis free-ware ImageJ (ImageJ, National Institute of Health, USA). The parameters used in the different functions were min_size = 5, max_size = 1000, linkrange = 2, disp = 20, net_filter = 10, duration_filter = 0.1, detect_filter = 0.1, median_step_filter = 3.

The species identification was achieved by comparing the traits extracted from species monocultures to each individual particle detected in videos of species in mixture using the Support Vector Machine algorithm (e1071 R-package, Meyer et al. 2014, function svm()). More specifically:

- to distinguish *Paramecium aurelia* from *Colpidium striatum*, we selected typical individuals in the monoculture with sufficient speed (net_speed > 10 and net_disp > 20) and we eliminated small *Paramecium* (argument: mean_minor > 45) and large *Colpidium* (argument: mean_minor < 35). Then we used the size traits (mean_minor + mean_major) in the model.
- to distinguish *Colpidium striatum* from *Euglena gracilis*, we selected the same *Colpidium* than above in the monoculture and the *Euglena* which were not moving to much, as they typically behave (net_speed < 7). We used all the traits (major_mean, major_sd, minor_mean, minor_sd, gross_speed_mean, gross_speed_sd, net_speed_mean, net_speed_sd, sd_turning_mean) in the model.

To check the probability of species assignation errors of the models, we applied them to the different monocultures. We then used the models on species in mixtures.

We did an additional visual check of the videos to avoid false negative of automated particle detection at low density for *Paramecium* and *Colpidium* (absence / presence records) and we counted the *Euglena* individuals visually on each video because their low speed was leading to systematic underestimations of the density.

## Supplementary Figures

**Figure S1.**
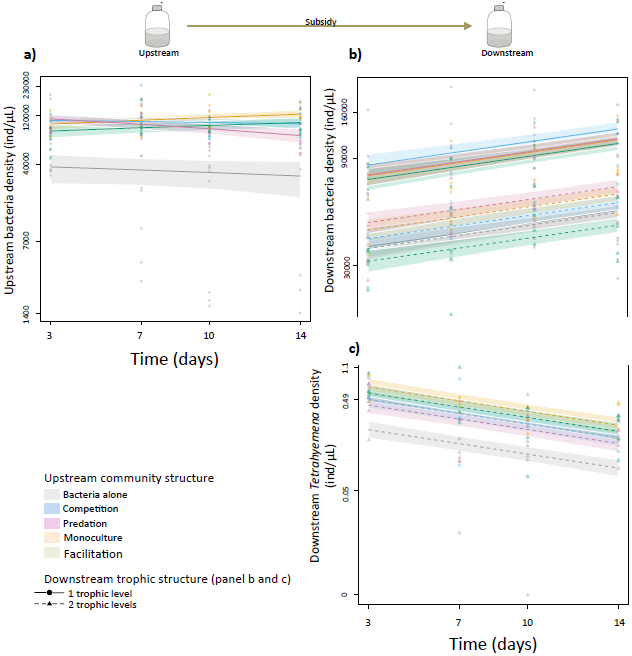
Experimental results with diluted resource. Effect of upstream community structure on upstream bacteria density (panel A), and on downstream bacteria (panel B) and *Tetrahymena* (panel C) densities in the one-trophic-level (full lines – *Tetrahymena* absent) and in the two-trophic-level (dashed lines – *Tetrahymena* present) communities. Points (*Tetrahymena* asbent) and triangles (Tetrahymena present) represent raw data. Full lines and dashed lines represent model predictions with 95% confidence intervals as shadings. Y-axes on all panels are on log-scale, but for clarity tick numbers represent raw densities.

**Figure S2.**
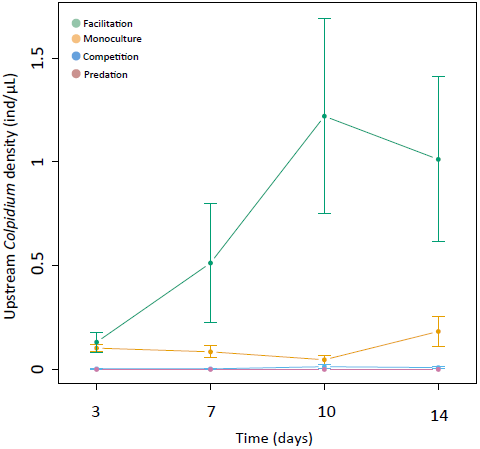
Upstream *Colpidium* density. Upstream *Colpidium* density as a function of community structure. Points represent MEAN ± SE.

## References

Altermatt, F. 2013. Diversity in riverine metacommunities: a network perspective. Aquatic Ecology 47:365–377.

Altermatt, F., E. A. Fronhofer, A. Garnier, A. Giometto, F. Hammes, J. Klecka, D. Legrand, E. Mächler, T. M. Massie, F. Pennekamp, M. Plebani, M. Pontarp, N. Schtickzelle, V. Thuillier, and O. L. Petchey. 2015. Big answers from small worlds: a user’s guide for protist microcosms as a model system in ecology and evolution. Methods in Ecology and Evolution 6:218–231.

Altermatt, F., S. Schreiber, and M. Holyoak. 2010. Interactive effects of disturbance and dispersal directionality on species richness and composition in metacommunities. Ecology 92:859–870.

Bourgeois, B., E. González, A. Vanasse, I. Aubin, and M. Poulin. 2016. Spatial processes structuring riparian plant communities in agroecosystems: implications for restoration. Ecological Applications 26:2103–2115.

Carrara, F., F. Altermatt, I. Rodriguez-Iturbe, and A. Rinaldo. 2012. Dendritic connectivity controls biodiversity patterns in experimental metacommunities. Proceedings of the National Academy of Sciences 109:5761–5766.

Carrara, F., A. Giometto, M. Seymour, A. Rinaldo, and F. Altermatt. 2014. Experimental evidence for strong stabilizing forces at high functional diversity of aquatic microbial communities. Ecology 96:1340–1350.

Dong, X., B. Li, F. He, Y. Gu, M. Sun, H. Zhang, L. Tan, W. Xiao, S. Liu, and Q. Cai. 2016. Flow directionality, mountain barriers and functional traits determine diatom metacommunity structuring of high mountain streams. Scientific Reports 6:24711.

Elkin, C. M., H. Possingham, A. E. Y. Michalakis, and E. D. L. DeAngelis. 2008. The role of landscape-dependent disturbance and dispersal in metapopulation persistence. The American Naturalist 172:563–575.

Fox, J., and S. Weisberg. 2011. An R companion to applied regression. Sage, Thousand Oaks, CA. URL: http://socserv.socsci.mcmaster.ca/jfox/Books/Companion.

Fronhofer, E. A., and F. Altermatt. in review. Classical metapopulation dynamics and eco-evolutionary feedbacks in dendritic networks. Ecography. bioRxiv: http://dx.doi.org/10.1101/033639.

Fronhofer, E. A., J. Klecka, C. J. Melián, and F. Altermatt. 2015a. Condition-dependent movement and dispersal in experimental metacommunities. Ecology Letters 18:954–963.

Fronhofer, E. A., T. Kropf, and F. Altermatt. 2015b. Density-dependent movement and the consequences of the Allee effect in the model organism Tetrahymena. Journal of Animal Ecology 84:712–722.

Fronhofer, E. A., N. Nitsche, and F. Altermatt. in press. Information use shapes the dynamics of range expansions into environmental gradients. Global Ecology and Biogeography. bioRxiv: http://dx.doi.org/10.1101/056002.

Gounand, I., E. Harvey, and F. Altermatt. in review. Subsidies mediate interactions between communities across space. Oikos.

Gounand, I., N. Mouquet, E. Canard, F. Guichard, Hauzy Céline, and D. Gravel. 2014. The paradox of enrichment in metaecosystems. The American Naturalist 184:752–763.

Gravel, D., F. Guichard, M. Loreau, and N. Mouquet. 2010a. Source and sink dynamics in meta-ecosystems. Ecology 91:2172–2184.

Gravel, D., N. Mouquet, M. Loreau, and F. Guichard. 2010b. Patch dynamics, persistence, and species coexistence in metaecosystems. The American Naturalist 176:289–302.

Harvey, E., I. Gounand, P. Ganesanandamoorthy, and F. Altermatt. 2016. Spatially cascading effect of perturbations in experimental meta-ecosystems. Proc. R. Soc. B 283:20161496.

Hauzy, C., F. D. Hulot, A. Gins, and M. Loreau. 2007. Intra- and interspecific density-dependent dispersal in an aquatic prey–predator system. Journal of Animal Ecology 76:552–558.

Kuglerová, L., R. Jansson, R. A. Sponseller, H. Laudon, and B. Malm-Renöfält. 2014. Local and regional processes determine plant species richness in a river-network metacommunity. Ecology 96:381–391.

Leibold, M. A., M. Holyoak, N. Mouquet, P. Amarasekare, J. M. Chase, M. F. Hoopes, R. D. Holt, J. B. Shurin, R. Law, D. Tilman, M. Loreau, and A. Gonzalez. 2004. The metacommunity concept: a framework for multi-scale community ecology. Ecology Letters 7:601–613.

Levine, J. M. 2003. A patch modeling approach to the community-level consequences of directional dispersal. Ecology 84:1215–1224.

Loreau, M., N. Mouquet, and R. D. Holt. 2003. Meta-ecosystems: a theoretical framework for a spatial ecosystem ecology. Ecology Letters 6:673–679.

Lutscher, F., E. McCauley, and M. A. Lewis. 2007. Spatial patterns and coexistence mechanisms in systems with unidirectional flow. Theoretical Population Biology 71:267–277.

Lutscher, F., E. Pachepsky, and M. Lewis. 2005. The effect of dispersal patterns on stream populations. SIAM Journal on Applied Mathematics 65:1305–1327.

Marleau, J. N., F. Guichard, and M. Loreau. 2015. Emergence of nutrient co-limitation through movement in stoichiometric meta-ecosystems. Ecology Letters 18:1163–1173.

Muneepeerakul, R., E. Bertuzzo, H. J. Lynch, W. F. Fagan, A. Rinaldo, and I. Rodriguez-Iturbe. 2008. Neutral metacommunity models predict fish diversity patterns in Mississippi–Missouri basin. Nature 453:220–222.

Nash, J. C. 1990. Compact Numerical Methods for Computers. Linear Algebra and Function Minimisation. Second edition. CRC press.

Pennekamp, F., and N. Schtickzelle. 2013. Implementing image analysis in laboratory-based experimental systems for ecology and evolution: a hands-on guide. Methods in Ecology and Evolution 4:483–492.

Pennekamp, F., N. Schtickzelle, and O. L. Petchey. 2015. BEMOVI, software for extracting behavior and morphology from videos, illustrated with analyses of microbes. Ecology and Evolution 5:2584–2595.

Pinheiro, J., D. Bates, S. DebRoy, D. Sarkar, and R Core Team. 2016. nlme: linear and nonlinear mixed effects model. URL: http://CRAN.R-project.org/package=nlme.

Polis, G. A., and S. D. Hurd. 1995. Extraordinarily high spider densities on islands: flow of energy from the marine to terrestrial food webs and the absence of predation. Proceedings of the National Academy of Sciences 92:4382–4386.

R Core Team. 2016. R: A language and environment for statistical computing. R Foundation for Statistical Computing, Vienna, Austria.

Salomon, Y., S. R. Connolly, and L. Bode. 2010. Effects of asymmetric dispersal on the coexistence of competing species. Ecology Letters 13:432–441.

Seymour, M., E. A. Fronhofer, and F. Altermatt. 2015. Dendritic network structure and dispersal affect temporal dynamics of diversity and species persistence. Oikos 124:908–916.

Sitters, J., C. L. Atkinson, N. Guelzow, P. Kelly, and L. L. Sullivan. 2015. Spatial stoichiometry: cross-ecosystem material flows and their impact on recipient ecosystems and organisms. Oikos 124:920–930.

Vannote, R. L., G. W. Minshall, K. W. Cummins, J. R. Sedell, and C. E. Cushing. 1980. The river continuum concept. Canadian Journal of Fisheries and Aquatic Sciences 37:130–137.

Venables, W. N., and B. D. Ripley. 2002. Modern applied statistics with S. Springer, New York.

Vitorino Júnior, O. B., R. Fernandes, C. S. Agostinho, and F. M. Pelicice. 2016. Riverine networks constrain β-diversity patterns among fish assemblages in a large Neotropical river. Freshwater Biology 61:1733–1745.

